# Sources of Variation in Fecal Haptoglobin in a Population of Wild Capuchin Monkeys (*Cebus imitator*)

**DOI:** 10.1101/2025.06.26.661772

**Authors:** Raquel Hernández-Rojas, Hadjira Hamou, Ronald López Navarro, Katharine M. Jack, Fernando A. Campos, Amanda D. Melin, James P. Higham

**Author notes:** **Correspondence:** Raquel Hernández-Rojas, University of Calgary, HMRB#202, 3330 Hospital Dr NW, Calgary, AB T2N 4N1.

## Abstract

Non-human primates support ecosystem function by enhancing forest regeneration through seed dispersal and other key ecological roles. Unfortunately, primate populations are declining, placing renewed emphasis on monitoring the health of wild populations. Non-invasive monitoring of reliable biomarkers of inflammation and immune activation allows researchers to assess individual health status without capturing or interfering with wild animals, but studies are limited by the availability of such biomarkers that are measurable from fecal and urine samples. In the present study, we aimed to validate the measurement of fecal haptoglobin, a biomarker of inflammation, in wild white-faced capuchin monkeys (*Cebus imitator*), and to evaluate the relationship between fecal haptoglobin concentrations and age, sex, dominance rank, circadian effects and environmental factors including temperature and rainfall. After analytically validating the measurement of fecal haptoglobin, our results did not demonstrate a relationship between haptoglobin concentrations and age, sex, dominance rank or circadian effects. However, we found significant influences of environmental conditions on fecal haptoglobin levels, with an increase and more variation observed during drier conditions, when the animals are typically under greater environmental stress. We conclude that haptoglobin measurement is feasible in wild white-faced capuchin monkeys, and its concentrations vary in our study population, reflecting seasonal patterns of inflammation that are consistent with changes to environmental stressors associated with lower access to food and water.

## Introduction

Non-human primates are distributed across multiple continents, supporting forest regeneration, contributing to ecosystem health, and playing an important role in the study of emerging diseases (Estrada et al., 2017; Galán-Acedo et al., 2019). Research on primate condition in the wild allows us to link variation in ecological and social pressures to changes in health and disease across the lifespan, topics that are challenging to study in captive conditions (Harrison & Van De Waal, 2022; Lopresti-Goodman & Villatoro-Sorto, 2022). Climate change has additionally brought global concern for the health and sustainability of primate populations (Carvalho et al., 2019). The world population of wild non-humans primates is decreasing due to habitat loss and trade in live primates (Estrada et al., 2017; Norconk et al., 2020), creating greater urgency of efforts to monitor the health of primates as a crucial component of their conservation (Bicca-Marques et al., 2022). Monitoring the health of wild primates can also aid in the early identification of zoonotic and anthroponotic diseases (Balansard et al., 2019). This highlights the importance of longitudinal studies that enable the monitoring of health over time, helping us to understand the fundamental biology of aging and variation in lifespan (Campos et al., 2024). Additionally, longitudinal studies allow researchers to examine the impact of seasonal or long-term environmental changes on nutrition and health, social behaviour, and the ecological roles of primates (Alberts & Altmann, 2012; Gonçalves et al., 2022).

At some field sites blood can be collected in order to take measurements of animal condition, but this requires direct manipulation of the animal. Such intervention can significantly impact the health of individuals due to the high levels of stress they may be subjected to, potentially leading to severe consequences, such as capture myopathy, which can be fatal (Breed et al., 2019). Capturing wild primates also has a negative impact on their normal behaviour, potentially influencing the validity of studies of natural behavior and life history, which has led to the search for new methods to collect information without disrupting their daily activity (Lopresti-Goodman & Villatoro-Sorto, 2022; Piel et al., 2022). Non-invasive sample collection generally causes minimal disruption to the normal behavior of the individual in the wild, and it lowers risks to researchers associated with sampling, particularly the potential transmission of zoonotic diseases transmitted through direct contact within animals (Behringer & Deschner, 2017; Smiley Evans et al., 2015). As part of these ongoing efforts, there is continued interest in expanding the number of non-invasive health biomarkers that are validated for use in wild and naturalistic primate populations (Lucore et al., 2022).

One potential biomarker of interest is haptoglobin, a glycoprotein whose levels in blood plasma changes during the acute phase of inflammatory responses (Hooijberg & Cray, 2023). Haptoglobin has three major subtypes: Hp1-1, Hp2-2 and Hp2-2, with Hp1-1 found in almost all animal species (Carter & Worwood, 2007; Lai et al., 2008). This protein can also be found in other body fluids across a variety of mammals (Wan et al., 2021). Blood concentrations of haptoglobin are considered a biomarker of inflammation and can be used in the monitoring of many systemic diseases (Naryzhny & Legina, 2021), while fecal levels of haptoglobin have been described as an important tool for predicting the presence of bowel lesions and the early diagnosis of colorectal cancer in humans, which are local affections limited to the gut (Chalkias et al., 2011; Shiotani et al., 2014). Haptoglobin levels increase in infectious diseases, such as parasitic, bacterial and viral infections (Quaye, 2008) and during subclinical infections, providing a valuable tool for detection in wildlife, particularly when agent-specific methods are impractical (Vicente et al., 2019). The levels of this glycoprotein can also increase during chronic diseases, including cardiovascular diseases and cancer, which has led to its growing use in human medical research (Cheng et al., 2024). In rhesus macaques (*Macaca mulatta*), lymph node extirpation and intestinal biopsy sampling lead to an increase in fecal excretion of haptoglobin (Higham et al., 2015). Despite the clinical importance of haptoglobin, the literature on non-human primates is limited, creating a need for studies that help to understand sources of variation of this biomarker in the wild.

Climatic seasonality has been documented to impact immune function in nonhuman primates. In baboons (*Papio* spp.), a seasonal immune rhythm is described, with an increase of markers of inflammation such as Interleukin-6 and C-reactive protein reported during months with the higher temperatures (McFarlane et al., 2012). Additionally, a study conducted in the tropical dry forest in Guanacaste, Costa Rica, concluded that mantled howler monkeys (*Alouatta palliata*) have more labile body temperatures compared to humans (Thompson et al., 2014), suggesting that at least some non-human primates are more sensitive to changes in temperature. Seasonality can also impact triggers of inflammation indirectly. For example, gastrointestinal parasite species richness is higher during the dry season in red howler monkeys (*Alouatta seniculus*), brown spider monkeys (*Ateles hybridus*), and variegated capuchins (*Cebus versicolor*) (Rondón et al., 2017), which may lead to increased gut inflammation.

In the present study, we investigated variation in fecal haptoglobin excretion in a population of wild capuchin monkeys from the tropical dry forest of the Santa Rosa Sector of the Área de Conservación Guanacaste (ACG) in Guanacaste, Costa Rica. Capuchin monkeys share several similarities with humans, including a relatively long lifespan in relation to body size, complex social behaviour, and an omnivorous generalist diet (Campos et al., 2024). These primates, specifically brown capuchins (*Sapajus apella*), have also been used to study important diseases such as Alzheimer’s (Diehl Rodriguez et al., 2024). The white-faced capuchin monkeys (*Cebus imitator*) present in the ACG experience a long dry season of approximately 5 months, with temperatures that can exceed 37°C, with low humidity and scarce precipitation (Campos & Fedigan, 2009), which impact the resource availability. The middle of the dry season is typically associated with low food abundance while water scarcity is an important stressor of the late dry season, especially as remaining water sources become small, heavily used and contaminated with feces of many animals (Hogan & Melin, 2018; Melin et al., 2014; Orkin et al., 2019). Capuchins in ACG show signatures of local genetic adaptation to the extreme dry season, including genes linked to water balance and kidney function (Orkin et al., 2021). As water sources become scarce, capuchin monkeys must drink from the few remaining watering holes, which are considered an important source of parasitic infections such as *Strongyloides*, and with fewer fruits available, they eat more insects, which could serve as intermediate host for gastrointestinal parasites such as cestodes and acanthocephalans (Henriquez et al., 2025). During the hottest hours of the day in the dry season, capuchins also travel shorter distances, likely to conserve energy and avoid overheating (Campos & Fedigan, 2009).

We analyzed haptoglobin concentrations from 185 fecal samples collected from 68 unique individuals over a period of 8 months, spanning the dry and wet seasons. Our first aim was to perform an **analytical validation** of the measurement of fecal haptoglobin in wild white-faced capuchin monkeys. To achieve this, we used a commercial direct sandwich enzyme-linked immunosorbent assay (ELISA), designed for haptoglobin detection in serum, which has been successfully applied to the analysis of fecal extracts in rhesus macaques (Higham et al., 2015). Additionally, we investigated the relationship between haptoglobin concentrations and a range of intrinsic, social, and environmental covariates including **age, sex**, **dominance rank**, **circadian effects, temperature**, and **rainfall**.

## Methods

### Ethics statement

This animal study was reviewed and approved by the Animal Care Committee (ACC) of the University of Calgary in Canada (AC19-0167), Tulane’s Institutional Animal Care and Use Committee (Protocol #2432) and by the Sistema Nacional de Áreas de Conservación (SINAC) and the Área de Conservación Guanacaste (ACG: R-SINAC-ACG-PI-059-2022/ ACG-PI-033-2023ACG-PI-011-2024/, and CONAGEBIO (R-013-2022-OT-CONAGEBIO/R-042-2023-OT-CONAGEBIO)/ R-021-2025-OT-CONAGEBIO) in Costa Rica. Fecal samples were imported to Canada under Canadian Food Inspection Agency (CFIA) permits A-2023-06194-1 and A-2022-05488-4.

### Study site and subjects

This study was conducted in the tropical dry forest of Sector Santa Rosa, ACG, Guanacaste, Costa Rica (10.836° latitude; −85.615° longitude), at an elevation of 300 meters above sea level, with two distinct seasons: a dry season and a wet season. The study population consisted of wild adult white-faced capuchin monkeys (*Cebus imitator*) from five social groups, including males and females, that have been followed and documented in ACG for more than 40 years.

### Sample collection

We collected fecal samples from 68 unique individuals (26 males and 42 females) from five different social groups. The age range was from 5 to 30 years old. A total of 185 samples (2.72 ∓ 1.41 per individual) were collected from October 2022 to June 2023, between 6:00 am to 5:00 pm. Trained field technicians collected the samples non-invasively and opportunistically after observing an individual defecating. Samples were collected from the ground or leaves and transferred into 2 ml cryovials. In the field, fecal samples were temporarily stored in a portable cooler until they arrived at the field facilities, where they were transferred to a cryogenic liquid nitrogen dewar before being shipped to the University of Calgary for analysis. All samples arrived frozen to the laboratory and were immediately stored at −80°C.

### Fecal analysis

At the University of Calgary, we undertook extractions by freeze-drying fecal samples overnight then pulverizing them. We added 1 ml of the extraction buffer ELISA kit to 20 mg of pulverized fecal sample, vortexed, and centrifuged the mixture. We then collected the supernatant and discarded the pellet.

We measured fecal haptoglobin using a commercial ELISA kit (Monkey Haptoglobin ELISA Kit, Catalog No. HAPT-3, Life Diagnostics, Inc.), following the manufacturer’s instructions. The kit is designed for haptoglobin detection in serum and has been successfully used for fecal extraction analysis in rhesus macaques (Higham et al., 2015). We read the absorbance at 450 nm using Synergy HTX multi-mode reader (Biotek, Ref: S1LFA) with the software Gen5 3.11 (BioTek Instruments, 2020).

The fecal extracts were measured undiluted; however, fecal extractions with results outside the linear portion of the standard curve were remeasured. A total of 141 samples did not need remeasurement. For 23 extracts with low concentrations, we freeze-dried them overnight and concentrated the sample by resuspending them in 0.25ml of the extraction buffer ELISA kit. For 21 extracts with high concentrations, we diluted them with the extraction buffer ELISA kit using higher dilution factors: 10 samples were diluted 1:4, 1 sample was diluted 1:8, 9 samples were diluted 1:16, and 1 sample was diluted 1:32. To determine the final concentration we incorporated the dilution or concentration factor. Samples with coefficient of variation (CV) greater than 10% were remeasured. The inter-assay variation was 7.98% for the high concentration quality control and 12.65% for the low concentration quality control. The final fecal haptoglobin concentration is expressed in ng/g of feces.

### Analytical validation

To select the best dilution factor for determining haptoglobin concentration in fecal extracts, we analyzed extractions from 9 different individuals with unknown haptoglobin concentrations. These extracts were serially diluted from 1:1 to 1:8 to identify which dilution fell within the linear portion of the standard curve. Subsequently, to assess assay performance across the tested concentration range, we conducted a parallelism test, by selecting 9 haptoglobin extracts with previously measured concentrations close to the maximum value of the standard curve. These extracts were also serially diluted from 1:1 to 1:8, and the trendline of each diluted sample was compared to that of the trendline of the standard curve.

### Environmental and social covariates

Temperature data were collected every half hour throughout the study period using an environmental meter (Kestrel, Model: 5000) located in a protected location at the research station, which is approximately in the center of the study groups’ home ranges. We later determined that the temperature data from the Kestrel meter were moderated by its location, resulting in lower maximum temperatures and higher minimum temperatures relative to conditions in the field. We therefore made use of a second temperature data source, a HOBO Weather Station (Onset Corp) with a S-THC-M002 temperature sensor protected by a solar radiation shield, for which data collection began in August 2023, after the period in which fecal samples were collected for this study. Using 15 months of subsequent overlapping simultaneous temperature recordings between the Kestrel and the HOBO instruments (from August 2023 to March 2025), we fit a linear model of HOBO temperature data as a function of Kestrel data, and we used this model to predict temperature values during the sampling period from the Kestrel data collected during that time. To measure rainfall, we used a Metric Rain Gauge (Cole-Parmer Instrument Company, Model: 03319-10), recording accumulated rainfall once every 24 hours.

### Statistical analysis

All statistical analyses were performed using R software, version 4.4.2 (R Core Team, 2024). We modeled log-transformed haptoglobin concentrations using a generalized additive mixed model (GAMM). We included smooth (non-linear) terms for the following continuous predictors: age in years (separately by sex), the average maximum temperature over the 15 days prior to and including the day of sample collection, the sum of rainfall over the 30 days prior to and including the day of sample collection, and the number of minutes into the day starting from midnight. We also included fixed effects of dominance rank (alpha vs. non-alpha), sex, and the interaction between rank and sex.

As random effects, we included individual ID, because each individual had multiple measurements, allowing each individual to have their own intercept, as well as the study group that the individual belonged to when the sample was collected. Study group was usually the same for all the samples of a given individual, but some individuals, especially males, were sampled as members of multiple different study groups.

To fit the models, we used the mgcv R package, version 1.9-1 (Wood, 2011), and for generating and visualizing the predictions and partial effects, we used the R packages marginaleffects, version 0.24.0 (Arel-Bundock et al., 2024), and gratia, version 0.10.0 (Simpson, 2024).

## Results

### Aim 1: Analytical validation

We found that the set of undiluted fecal extracts fell within the linear portion of the standard curve, with a higher distribution around the lower end of the standard curve. For the fecal extracts diluted using the factors 1:2, 1:4, and 1:8, we found undetectably low results of 20%, 40%, and 60%, respectively, for each dilution factor. For this reason, we ran all samples undiluted in this study. However, a total of 23 of the samples had undetectably low levels of haptoglobin and needed to be concentrated, 10 of them collected on rainy days. Additionally, 21 samples had undetectably high levels of haptoglobin and required further dilution. According to the date of sampling, all those samples were collected on dry days with scarce rainfall. In the parallelism test, we found that the trendlines of the samples exhibited parallel behaviour to the trendline of the standard curve, indicating consistent assay performance across different sample concentrations.

### Aim 2: Intrinsic, social, and environmental influences on haptoglobin

Fecal haptoglobin concentrations do not exhibit a significant relationship with age, and do not differ by sex, dominance status of the monkeys or circadian effects. (**Table 1**), as assessed using GAMM (**Figure 1**). There were significant effects of environmental conditions on fecal haptoglobin concentrations, which were higher during drier conditions, where the rainfall is low over the previous 30 days (**Table 1**).

**Figure 1.**
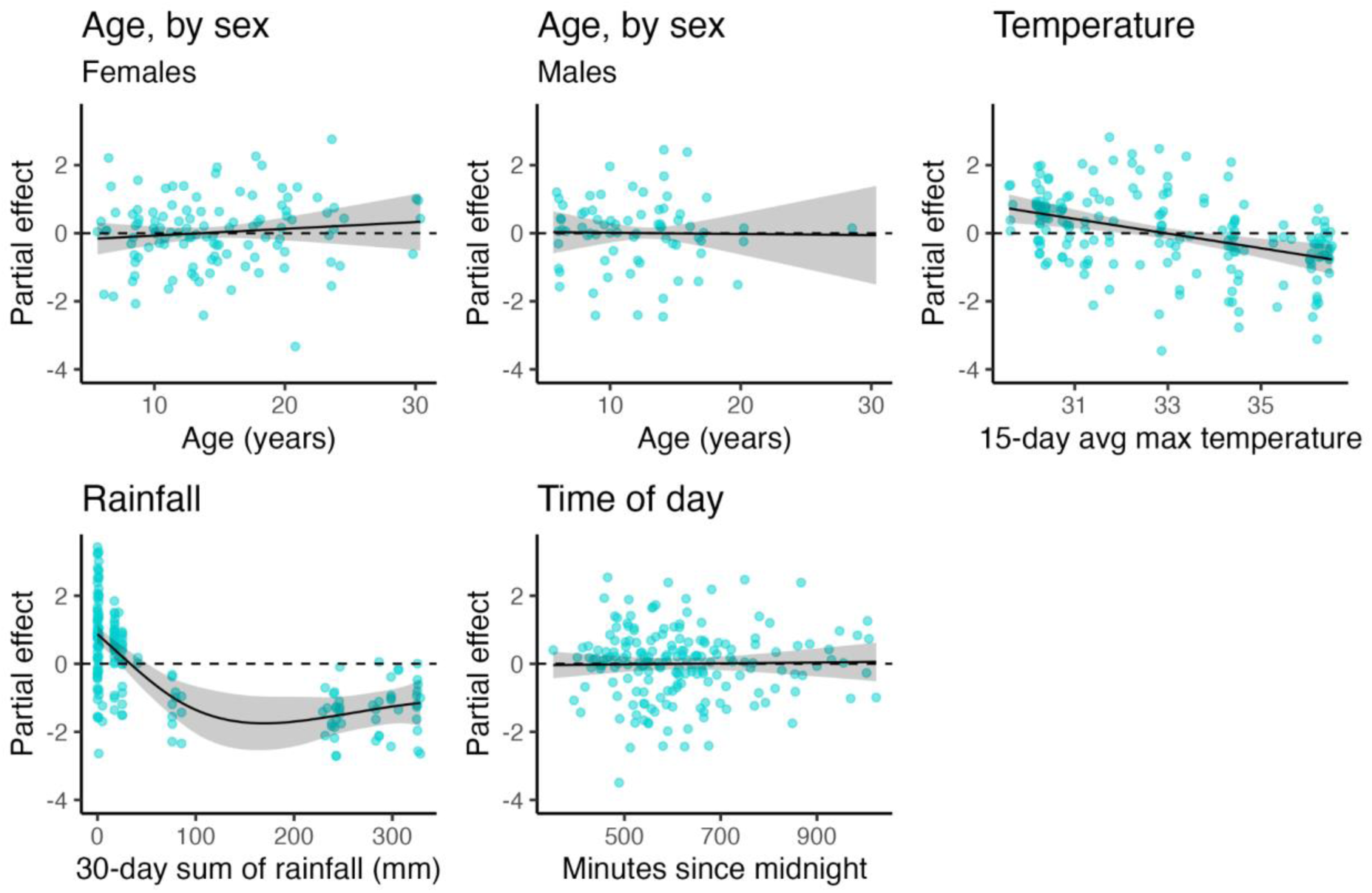
Partial effects and partial residuals of age (by sex), temperature, rainfall, and time of day on fecal haptoglobin concentration (ng/g feces) in wild white-faced capuchin monkeys. While no age, sex, or time-of-day effects were evident, haptoglobin concentrations were lower following rainy periods and were inversely related to temperature.

**Table 1.** Summary of parametric and smooth term results from the generalized additive mixed model.

| Parameter | Coefficient | SE | 95% CI | t / F | df | df (error) | p |
| --- | --- | --- | --- | --- | --- | --- | --- |
| **Parametric terms** |  |  |  |  |  |  |  |
| (Intercept) | 5.74 | 0.47 | ( 4.82, 6.67) | 12.28 |  | 147.53 | < .001 |
| rank [Sub] | -0.16 | 0.47 | (-1.09, 0.76) | -0.35 |  | 147.53 | 0.725 |
| sexMale | 0.22 | 0.57 | (-0.91, 1.36) | 0.38 |  | 147.53 | 0.701 |
| rank [Sub] ×<br>sexMale | -0.13 | 0.62 | (-1.35, 1.10) | -0.21 |  | 147.53 | 0.836 |
| **Smooth terms** |  |  |  |  |  |  |  |
| Smooth term<br>(age) ×<br>sexFemale |  |  |  | 0.59 | 1.00 |  | 0.445 |
| Smooth term<br>(age) ×<br>sexMale |  |  |  | 6.88e-03 | 1.00 |  | 0.934 |
| Smooth term<br>(temperature) |  |  |  | 13.81 | 1.00 |  | < .001 |
| Smooth term<br>(rainfall) |  |  |  | 18.07 | 2.89 |  | < .001 |
| Smooth term<br>(time of day) |  |  |  | 0.04 | 1.00 |  | 0.842 |
| Smooth term<br>(monkey id) |  |  |  | 0.65 | 24.45 |  | 0.006 |
| Smooth term<br>(study group) |  |  |  | 2.52 | 2.13 |  | 0.073 |

## Discussion

In this study we validated the measurement of fecal haptoglobin in wild white-faced capuchin monkeys by ELISA. As part of the standardization of fecal haptoglobin measurement in capuchin monkeys, our results suggest that samples collected during extreme climatic conditions may need to be manipulated via dilution for those collected during dry days with scarce rainfall, or via concentration for samples collected during rainy days, to enable reliable measurement and downstream analyses. This approach aims to reduce unnecessary consumption of samples and resources. With the parallelism test, we demonstrated that the standard curve exhibits similar behaviour to dilutions of the fecal extracts from capuchin monkeys for all tested samples.

The study of haptoglobin as a nonspecific biomarker of inflammation in platyrrhine primates is limited, and there has been no exploration of its relation to age, sex, dominance rank, circadian effects or climate variability. In humans, differences in serum haptoglobin concentrations have been reported according to sex and age. In neonates haptoglobin levels are extremely low, often making them undetectable (Jacob, 2016). A study in children aged 1 to 12 years old found that serum haptoglobin concentrations increase with age in those with higher densities of the malaria parasite (Fowkes et al., 2006). It has also been reported that women and older individuals tend to present higher levels; however, serum haptoglobin concentration can also be affected by the proportions of haptoglobin subtypes, which could explain why sex and age effects are not observed in other studies when some subtypes are overrepresented (Kasvosve et al., 2000; Lei et al., 2023). In this study, we detected no difference in haptoglobin with respect to the age, sex, dominance rank, circadian effects or dominance status of the monkeys. However, due to the scarce research available on this biomarker in platyrrhine primates, information about haptoglobin subtypes and their proportions in this population is not available.

We found significant influences of environmental conditions on fecal haptoglobin concentrations, which were higher during drier conditions. The tropical dry forest environment of ACG presents a challenge during the dry season for the capuchin monkeys, in terms of food availability, water scarcity and parasitism, which have the potential to create chronic physiological stress for their bodies (Henriquez et al., 2025; Melin et al., 2014; Orkin et al., 2019). A commonality of these variables is they all impact the intestinal health of primates and are exacerbated in the dry season. When animals do not have enough food available, it leads to a decrease in the intestinal transit speed, increases epithelial permeability in the gut, causes an imbalance in the intestinal microbiota, and consequently leads to intestinal inflammation (Genton et al., 2015). A separate study of our population demonstrated that seasonality influences the composition and function of their gut microbiome. When the fruit availability was scarce, microorganisms such as *Campylobacter, Enterococcus, Helicobacter, Haemophilis, Pseudomonas,* and *Streptococcus,* which are associated with human dysbiosis, ill health, and irritable bowel syndrome, increased (Orkin et al., 2019). Furthermore, the scarcity of water has a direct impact on intestinal homeostasis, leading to dysbiosis of the gut microbiome and reduction in the number of certain immune cells, which decreases the ability to eliminate microorganisms (Sato et al., 2024).

Previous research on capuchins in ACG has also found higher rates of infection by some gastrointestinal parasites during the dry season, when animals are more exposed to parasitic infections with *Strongyloides*, cestodes, and acanthocephalans (Henriquez et al., 2024; Henriquez et al., 2025). These infections can cause diarrhea and weight loss due to improper nutrient absorption. Consequently, this affects gut health and leads to inflammation (Dib et al., 2023). Thus, the rise in fecal haptoglobin levels we found in association with drier conditions could be explained by the damage and onset inflammation that the parasites can cause to the gut of the host. Another study in wild black capuchin monkeys demonstrated that parasite loads decreased when the animals had higher food availability, which could indicate that nutritional status affects the parasite dynamics in non-human primates (Agostini et al., 2017). This may be relevant for our population, as haptoglobin levels are lower in the months when food availability is higher (Melin et al., 2014).

In sum, we found that fecal haptoglobin measurement is viable in wild white-faced capuchin monkeys, and that concentrations of this biomarker vary in response to seasonally driven climatic variation in our study population. Future studies integrating this biomarker of health and inflammation could combine data on the gut microbiome and other measures of animal condition as part of integrated studies addressing the sources of variation in individual health across the lifespan.

## Supporting information

Anonymized data

## Abbreviations

ACG: Área de Conservación Guanacaste
ELISA: enzyme-linked immunosorbent assay
GAMM: generalized additive mixed model

## Acknowledgements

We thank Roger Blanco, María Marta, Staff and Administration of ACG at Sector Santa Rosa. Linda M Fedigan, Saúl Chéves Hernández, Danielka Rugama Taylor, Wendy Téllez Arias, Philippine Delga, Lindy Wolhuter, Cielo De La Rosa Meza, Ann-Kathrin Pohle, Peyton Schmidli, Suheidy Romero Morales, and all members of the Santa Rosa field team. We thank Patricia Ströher for administrative support.

## Notes

### Competing Interest Statement

The authors have declared no competing interest.

### Funding

- Natural Sciences and Engineering Research Council of Canada: NSERC; RGPIN-2023-04399, DGECR-2023-00272, RGPIN-2017-03782
- Canada Research Chairs program: 950-231257
- National Institutes of Health Aging Research In Animals: R61-AG078529-01
- National Science Foundation: NSF BCS 2127375, NSF SES 1919305

